# Synaptotagmin-9 In Mouse Retina

**DOI:** 10.1101/2023.06.27.546758

**Authors:** Chris S. Mesnard, Cassandra L. Hays, Lou E. Townsend, Cody L. Barta, Channabasavaiah B. Gurumurthy, Wallace B. Thoreson

**Affiliations:** Truhlsen Eye Institute and Department of Ophthalmology and Visual Sciences, University of Nebraska Medical Center, Omaha, NE 68106, USA; Pharmacology and Experimental Neuroscience, University of Nebraska Medical Center, Omaha, NE 68106, USA; Genetics, Cell Biology and Anatomy, University of Nebraska Medical Center, Omaha, NE 68106, USA; Department of Medical Education, Creighton University, Omaha, NE 68178

**Keywords:** Synaptotagmin, electroretinogram, rod photoreceptor cell, retina

## Abstract

Synaptotagmin-9 (Syt9) is a Ca^2+^ sensor mediating fast synaptic release expressed in various parts of the brain. The presence and role of Syt9 in retina is unknown. We found evidence for Syt9 expression throughout the retina and created mice to conditionally eliminate Syt9 in a cre-dependent manner. We crossed Syt9^fl/fl^ mice with Rho-iCre, HRGP-Cre, and CMV-cre mice to generate mice in which Syt9 was eliminated from rods (rod^Syt9CKO^), cones (cone^Syt9CKO^), or whole animals (CMV^Syt9^). CMV^Syt9^ mice showed an increase in scotopic electroretinogram (ERG) b-waves evoked by bright flashes with no change in a-waves. Cone-driven photopic ERG b-waves were not significantly different in CMV^Syt9^ knockout mice and selective elimination of Syt9 from cones had no effect on ERGs. However, selective elimination from rods decreased scotopic and photopic b-waves as well as oscillatory potentials. These changes occurred only with bright flashes where cone responses contribute. Synaptic release was measured in individual rods by recording anion currents activated by glutamate binding to presynaptic glutamate transporters. Loss of Syt9 from rods had no effect on spontaneous or depolarization-evoked release. Our data show that Syt9 is acts at multiple sites in the retina and suggest that it may play a role in regulating transmission of cone signals by rods.

## Introduction

Synaptotagmins are a family of Ca^2+^-sensing proteins that facilitate synaptic exocytosis. Synaptotagmins -1, -2, and -9 serve as Ca^2+^ sensors that trigger fast synchronous release in CNS neurons (Geppert et al., 1994; Pang et al., 2006; Xu et al., 2007). The less-studied Synaptotagmin-9 (Syt9) shares considerable sequence homology with Syt1 and Syt2 (Rickman et al., 2004). Due to a nomenclature issue in the earlier literature, Syt9 was sometimes referred to as Syt5 and vice versa (Fukuda and Sagi-Eisenberg, 2008), so one must use care when reviewing the literature. For these experiments, we used Syt9 NCBI gene ID 60510 (491 amino acids). In the brain, Syt9 is expressed primarily in the hypothalamus, accumbens, striatum, and olfactory bulb (Mittelsteadt et al., 2009; Xu et al., 2007) (http://mouse.brain-map.org). Syt9 has been found to mediate fast, synchronous release in these areas, albeit with slower kinetics than Syt1 or Syt2 (Kochubey et al., 2016; Xu et al., 2007), along with a lower Ca^2+^ affinity (Hui et al., 2005). Eliminating Syt9 from striatal neurons was abolished fast synchronous release of GABA (Xu et al., 2007) but a different study in cultured neurons (hippocampal, cortical, and striatal) found that deletion of Syt9 had no effect on evoked inhibitory currents (Seibert et al., 2022). However, this same study showed that in Syt1 KO neurons, overexpressing Syt9 to levels to match Syt1 expression (∼25-fold) was able to rescue loss of synchronous release. Elimination of Syt9 in cultured striatal neurons also resulted in lower spontaneous release of GABA (Seibert et al., 2022). Syt9 is present on both small synaptic and large dense core vesicles (DCVs) of the mouse brain (Fukuda, 2006), and has been found to regulate catecholamine secretion (along with Syt1) from DCVs in cultured PC12 cells (Fukuda et al., 2002; Lynch and Martin, 2007). In the gonadotrophs of female mice, Syt9 has been shown to regulate the release of follicle-stimulating hormone (FSH) from dense core vesicles (Roper et al., 2015).

Nothing is known about possible roles of Syt9 in retina. Consistent with evidence for Syt9 genes in zebrafish retina (Henry et al., 2022), we show evidence for Syt9 in multiple retinal neurons. Selective elimination of Syt9 from rods caused a reduction in scotopic b-waves and oscillatory potentials, while global elimination of Syt9 increased b-waves, suggesting actions at multiple sites in the retina.

## Materials and methods

### Mice

We used control C57Bl6J and mutant mice aged 4–8 weeks for these experiments. Creation of HRGP-Cre and Rho-iCre mice have been described previously (Le et al., 2004; Li et al., 2005). Creation of Syt9^fl^ mice are described elsewhere (Quadros et al., 2017). Syt9^fl^ mice were originally created using C57Bl6N mice and then back-crossed to C7Bl6J mice. Yun Le (Univ. of Oklahoma) generously provided HRGP-Cre mice. Rho-iCre mice were obtained from Jackson Laboratories, Bar Harbor, ME (B6.Cg-Pde6b+ Tg(Rho-iCre)1Ck/Boc; RRID: 015850). Rho-iCre and HRGP-Cre mice selectively express cre-recombinase in rods and cones, respectively (Jin et al., 2020; Le et al., 2004; Li et al., 2005). CMV-Cre mice obtained from Jackson Laboratories (B6.C-Tg(CMV-cre)1Cgn/J) produce ubiquitous expression of cre-recombinase under control of a minimal human cytomegalovirus promoter.

Euthanasia was conducted in accordance with AVMA Guidelines for the Euthanasia of Animals by CO2 asphyxiation followed by cervical dislocation. Animal care and handling protocols were approved by the University of Nebraska Medical Center Institutional Animal Care and Use Committee.

### RNAscope

Eyes were fixed in either 4% paraformaldehyde (PFA) or 10% neutral buffered formalin (NBF) for 48 hours, transferred to 70% ethanol, and processed for paraffin embedding (UNMC Tissue Sciences Facility). The following steps were performed using the RNAscope™ 2.5 HD Reagent Kit (RED) along with the Quick Guide for FFPE Tissues (Advanced Cell Diagnostics, Newark, CA; Cat No: 322350). Five-micron thick retina slices were baked at 60° C, followed by deparaffinization in a series of xylene and 100% ethanol baths at room temperature. Slides were set to air dry for 5 minutes to prepare for pretreatment. Pretreatment of slides included direct application of hydrogen peroxide to tissue sections for 10 minutes followed by consecutive distilled water washes. Heat-induced antigen retrieval was completed by submerging sections into Target Retrieval solution at 95-100° C, sustained by hot water bath. Prior to in situ hybridization, tissue was digested using Protease Plus enzyme solution at 40°C in a humidity chamber.

In situ hybridization for Syt9 RNA was performed using probes targeting Syt9 (Advanced Cell Diagnostics, Newark, CA; Lot #:20293B). Probe Mm-Syt 9 (Cat No. 845291 from ACD Bio) targets the base pair region 636 - 1595 of the Syt9 transcript (NCBI accession number NM_021889.4). This target region includes a portion of exon 2; the entirety of exons 3, 4, and 5; and a portion of exon 6. Target amplification and color development was done using the 2.5 HD Red Detection sub-kit. Visualization was achieved utilizing Fast Red dye and sections were counterstained with Hematoxylin QS (Vector Laboratories, Burlingame, CA). Images were captured using SPOT Basic software (SPOT Imaging, Diagnostic Instruments Inc.) on a Spot Idea color camera (Diagnostics Instruments, Model 28.2) and Leitz Diaplan upright microscope with 40X objective.

### Electroretinography

ERGs were recorded in vivo using a UTAS Sunburst ganzfeld illuminator (LKC, Gaithersburg, MD, LKC-UTAS-SB). Mice were dark-adapted for ∼12 h prior to experiments and then anaesthetized via intra-peritoneal injection with a ketamine/xylazine drug cocktail (100 mg/kg ketamine, 10 mg/kg xylazine). Core temperature of the mouse was maintained at 37 °C with a heat pad. Tropicamide and proparacaine ophthalmic solution (0.5%) were administered topically to the left eye before the mouse was secured to the platform and a silver/silver chloride wire ring recording electrode was centered on the left cornea. Subcutaneous ground and reference electrodes were placed at the base of the tail and under the scalp, respectively. ERG a-waves provide a measure of photoreceptor responses and were measured from baseline to the bottom of the downward going negative potential. ERG b-waves reflect responses of second-order ON-type bipolar cells and were measured from the trough of the initial negative deflection of the a-wave to the peak of the positive-going b-wave. Measurements in dark-adapted (scotopic) conditions involved flashes of increasing intensity: 51 dB, −45 dB, −39 dB, −33 dB, −27 dB, −21 dB, −15 dB, −9 dB, −3 dB, and +5 dB. The intensity at 0 dB = 2.5 cd·s/m^2^. Ten flashes were presented at each intensity, separated by 10 s for steps 1–9 and 20 s between flashes at the highest intensity. Light-adapted (photopic) protocols were performed after background adaptation for 10 min with green light (40 cd/m2) and conducted with the same background. We tested 6 intensities (−6, −3, 0, 4, 7, and 13 dB) with 25 flashes at each intensity separated by 3 s. Oscillatory potentials were extracted from flash ERGs by bandpass filtering the entire trace to remove frequency components below 70 Hz and above 280 Hz using an 8-pole Bessel filter in Clampfit (Axon Instruments, Molecular Devices).

### Whole cell recordings

Whole cell recordings from rods were performed using a flatmount preparation of isolated retina. After enucleation, each eye was immediately placed in Ames’ medium (US Biological; RRID:SCR_013653) bubbled with 95% O_2_/5% CO_2_. The cornea was punctured with a scalpel and the anterior segment removed. The retina was isolated after cutting optic nerve attachments. After making four fine cuts at opposite poles, the retina was flattened onto a glass slide in the perfusion chamber with photoreceptors facing up. The retina was anchored in place with a brain slice harp (Warner Instruments, cat. no. 64-0250). The perfusion chamber was placed on an upright fixed-stage microscope (Nikon E600FN) equipped with a 60x water-immersion, long-working distance objective (1.0 NA). Flatmount preparations were superfused with room temperature Ames solution bubbled with 95%/5%CO2 at ∼1 mL /min. Outer segments were removed by gentle suction using a patch pipette to expose inner segments for recording.

Patch recording electrodes were pulled on a Narishige (Amityville, NY) PP-830 vertical puller using borosilicate glass pipettes (1.2-mm outer diameter, 0.9-inner diameter with internal filament; World Precision Instruments, Sarasota, FL). Pipettes had tip diameters of 1–2 *μ*m and resistances of 10– 15 M*Ω*. Rod inner segments and cell bodies were identified in flatmount retina and targeted with positive pressure using recording electrodes mounted on Huxley-Wall or motorized micromanipulators (Sutter Instruments). Cones were distinguished from rods by their larger membrane capacitance and much larger Ca^2+^ currents.

Rod ribbons are surrounded by the glutamate transporter EAAT5 and glutamate reuptake into rods by EAAT5 activates a large, anion conductance (Arriza et al., 1997; Eliasof et al., 1998; Grant and Werblin, 1996; Grassmeyer et al., 2019; Hasegawa et al., 2006; Picaud et al., 1995; Schneider et al., 2014; Thoreson and Chhunchha, 2023). I_A(Glu)_ is activated during glutamate re-uptake but is thermodynamically uncoupled from the transport process (Machtens et al., 2015). To enhance I, Cl^-^ in the patch pipette was replaced with a more permeable anion, thiocyanate (Eliasof and Jahr, 1996). The intracellular pipette solution for these experiments contained (in mM): 120 KSCN, 10 TEA-Cl, 10 HEPES, 1 CaCl2, 1 MgCl2, 0.5 Na-GTP, 5 Mg-ATP, 5 EGTA, 5 phospho-creatine, pH 7.2.

Rod I_A(Glu)_ recordings were performed in whole-cell voltage clamp using an Axopatch 200B amplifier (Molecular Devices) and signals were digitized with a DigiData 1550 (Molecular Devices). Data acquisition and analysis were performed using pClamp 10 software (Molecular Devices). Currents were acquired at 10 kHz and filtered at 2 kHz. Passive membrane resistance was subtracted from I_A(Glu)_ using P/6 subtraction. Membrane capacitance, membrane resistance, and access resistance values for rods using the KSCN solution averaged 3.0 ± 0.4 pF, 3.4± 2.0 GΩ, and 55 ± 9.5 MΩ (*n* = 9), respectively. Voltages were not corrected for a liquid junction potential of 3.9 mV. Chemical reagents were obtained from Sigma-Aldrich unless otherwise indicated.

### Statistical analysis

Statistical analysis and data visualization were done using ClampFit 10 and GraphPad Prism 9 software. Roughly equal numbers of male and female mice were used for these experiments. For ERG measurements, we analyzed the sample for each condition using Dunnett’s multiple comparisons test with one-way ANOVA. For comparing spontaneous release rates, we used nested t-tests to compare samples from cells where multiple measurements were made. Data values in the text are reported as mean ± SD. Errors bars for whole cell measurements in Fig. 7 show 95% confidence intervals.

## Results

### Localization of Syt9 mRNA in the retina

We used RNAscope techniques to localize sites of Syt9 mRNA in the retina. As illustrated in Fig. 1A-D, we first studied tissue fixed with 4% PFA. In control C56Bl6J mice, labeling for mRNA (red puncta) could be seen in rod photoreceptors (arrows, Fig. 1A), inner retinal neurons (Fig. 1A, B), and an occasional retinal ganglion cell (Fig. 1C). Rods showing RNAscope labeling for Syt9 are noted by arrows in the magnified section of Fig. 1B. Labeling for Syt9 mRNA in photoreceptor inner segments was eliminated in rod-specific conditional Syt9 knockout mice (Rod^Syt9CKO^, Fig. 1C-D). In retinas fixed with 10% NBF, mRNA labeling was diminished but we still saw labeling in photoreceptors and inner retinal neurons (arrows, Fig. 1F). Labeling was abolished entirely in a whole animal knockout of Syt9 (CMV^Syt9^) created by crossing floxed Syt9 mice with CMV-Cre mice that express cre-recombinase constitutively (Fig. 1G-H). Retinal thickness measured near the center of fixed retinas did not vary significantly among the three genotypes (WT: 220.4 + 15.2 μm, SD, n=5 mice; CMV^Syt9^, 213.5 + 15 μm, n=5 mice; Rod^Syt9CKO^, 221.5 + 25.4 μm, n=3 mice; P=0.735, 1-way ANOVA).

**Fig. 1.**
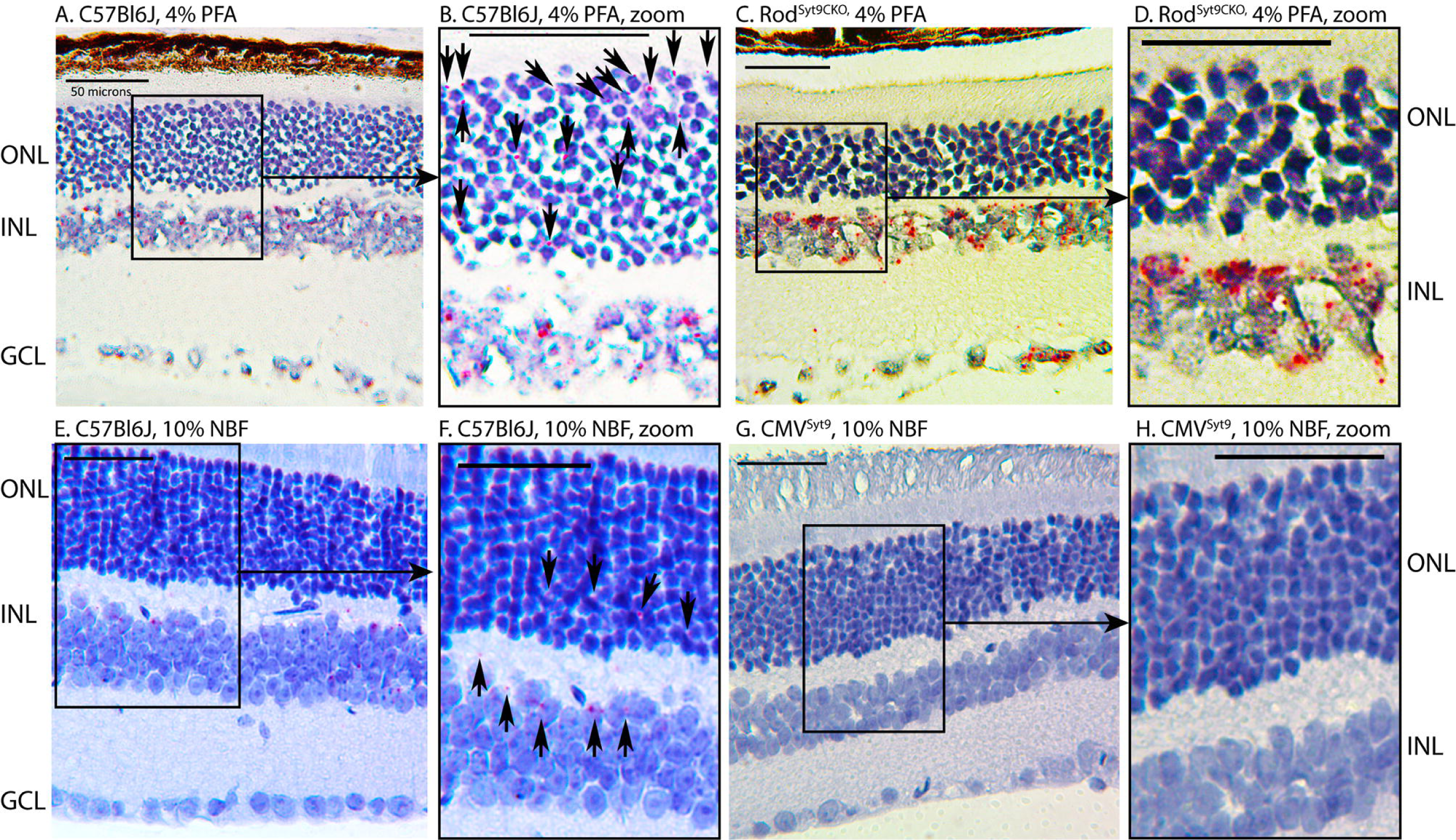
Syt9 mRNA visualized using RNAscope techniques (red puncta) was seen in the outer nuclear layer (ONL), inner nuclear layer (INL), and ganglion cell layer (GCL). Panels A-D show tissue fixed with 4% paraformaldehyde (PFA). Panels E-H show tissue fixed with 10% neutral buffered formalin (NBF). Panel A shows a retinal section of control C57Bl6J retina fixed with 4% PFA. Panel B shows a magnified region of the same section. Syt9 labeling (red puncta) in rods is indicated by the arrows in B. Panel C and magnified region in D show that labeling for Syt9 mRNA was eliminated from rods in rod-specific conditional Syt9 knockout mice (Rod^Syt9CKO^), but remained in inner retinal neurons and ganglion cells. Panel E shows a different control C57Bl6J retina fixed with 10% NBF. Labeling of neurons in the ONL and INL (F; arrows) was also seen with this fixative, but fewer cells were labeled. Labeling was abolished altogether in a whole animal knockout of Syt9 created by crossing floxed Syt9 mice with CMV-cre mice that express cre-recombinase constitutively (CMV^Syt9^).

### ERG effects of global elimination of Syt9

To study the role of Syt9 in synaptic transmission from photoreceptors in shaping bipolar cell light responses, we measured ERGs in vivo from anesthetized mice. Fig. 2A shows representative ERG responses evoked by a high intensity 20 ms flash applied under scotopic conditions to dark-adapted control mice (black trace) and mice in which Syt9 had been globally eliminated (CMV^Syt9^; red trace). The negative a-wave provides a measure of photoreceptor responses, while the positive b-wave is a measure of second-order ON-type bipolar cell responses. Fig. 2B and C display scotopic a- and b-wave amplitudes, respectively, as a function of increasing flash intensity. Example waveforms for photopic b-waves of light-adapted control (black) and CMV^Syt9^ (red) mice are shown in Fig. 2D. Fig. 2E plots the amplitude of photopic b-wave responses with increasing light intensity. Global elimination of Syt9 increased scotopic b-waves evoked by high intensity flashes by up to 37% (Fig. 2C), while scotopic a-waves (Fig. 2B) and photopic b-waves (Fig. 2E) were not significantly different.

**Fig. 2.**
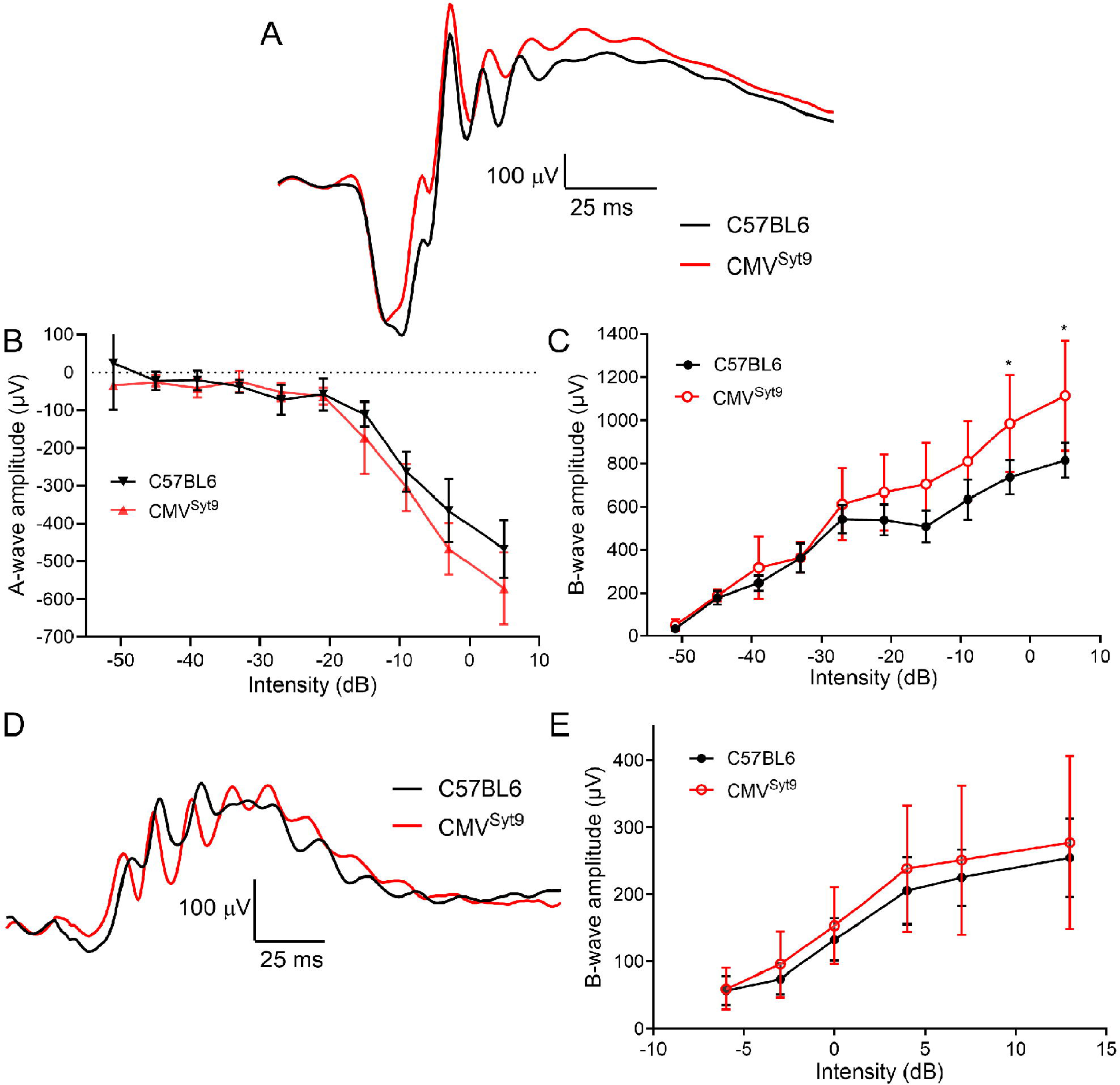
Global elimination of Syt9 increased scotopic b-wave responses at the highest flash intensities. (A). Representative scotopic ERG waveforms evoked from a high intensity flash (5dB) for control (n=6) and CMV^Syt9^ (n=7) mice are shown in black and red traces, respectively. (B). Scotopic a-wave amplitude as a function of flash intensity. (C). Plot of scotopic b-wave amplitude as a function of intensity. (D). Example photopic waveforms evoked from a high intensity flash (13 dB). (E). Plot of photopic b-wave amplitude as a function of intensity. *, P<0.01. Error bars show + S.D.

### ERG effects of selectively eliminating Syt9 from cones

Fig. 3 displays measurements from ERGs of control mice and Cone^Syt9CKO^ mice in which Syt9 was selectively eliminated from cones. Fig. 3A shows example scotopic ERG waveforms from dark-adapted control and Cone^Syt9CKO^ mice (black and red traces). Scotopic a- and b-wave amplitudes are plotted in Fig. 3B and C, respectively. Eliminating Syt9 from cones did not alter scotopic a- (Fig. 3B) or b-waves (Fig. 3C) significantly. Fig. 4D shows representative photopic ERG waveforms in light-adapted control and Cone^Syt9CKO^ mice. Photopic b-waves from Cone^Syt9CKO^ mice did not differ significantly from control mice (Fig. 3E).

**Fig. 3.**
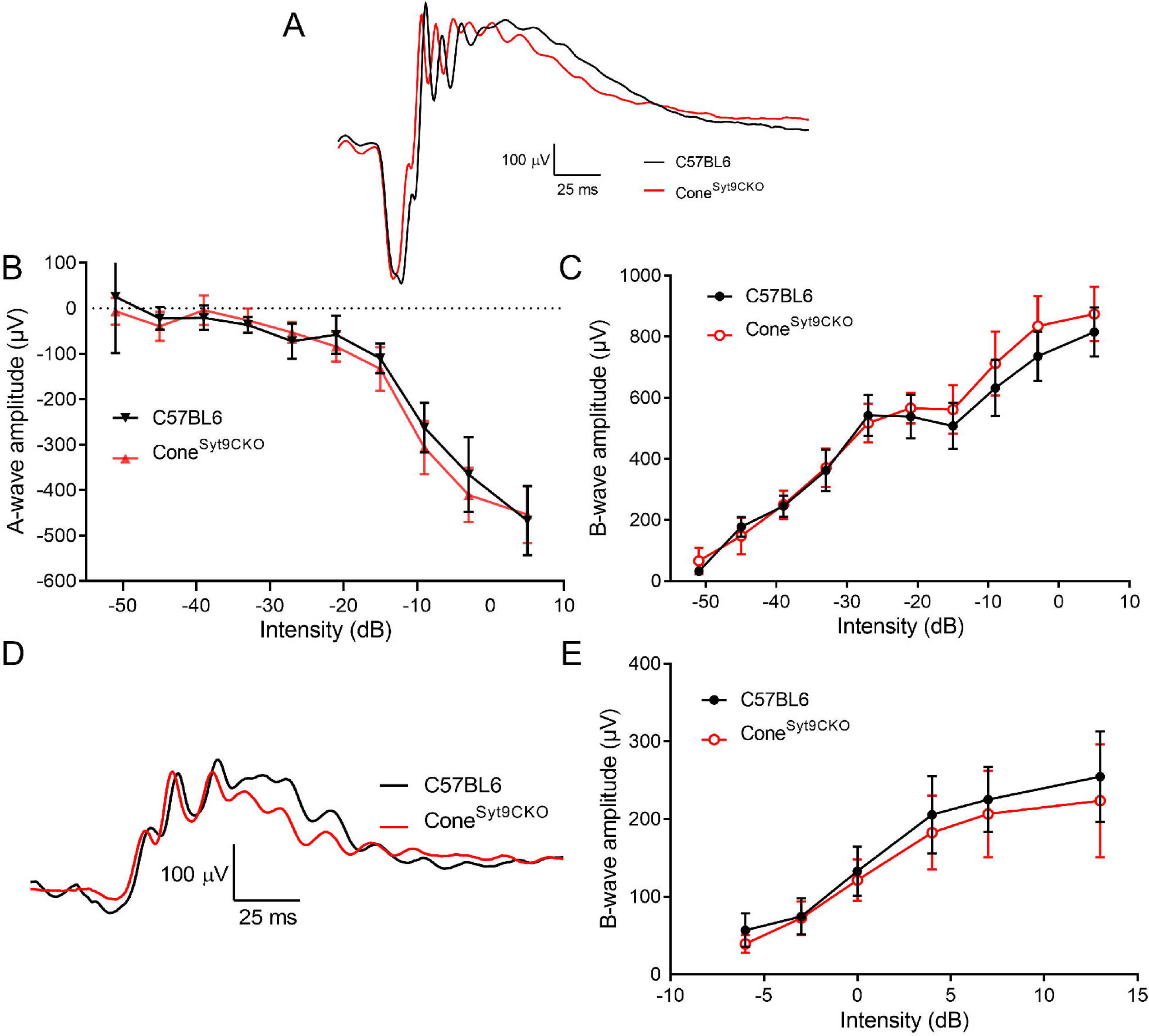
Elimination of Syt9 selectively from cones had no effect on scotopic or photopic ERG. (A). Representative scotopic ERG waveforms evoked from a high intensity flash (5 dB) for control (n=6) and Cone^Syt9CKO^ (n=7) mice are shown in black and red traces, respectively. (B). Scotopic a-wave amplitude as a function of flash intensity. (C). Plot of scotopic b-wave amplitude as a function of intensity. (D). Example photopic waveforms evoked from a high intensity flash (13 dB). (E). Plot of photopic b-wave amplitude as a function of intensity. Error bars show + S.D.

**Fig. 4.**
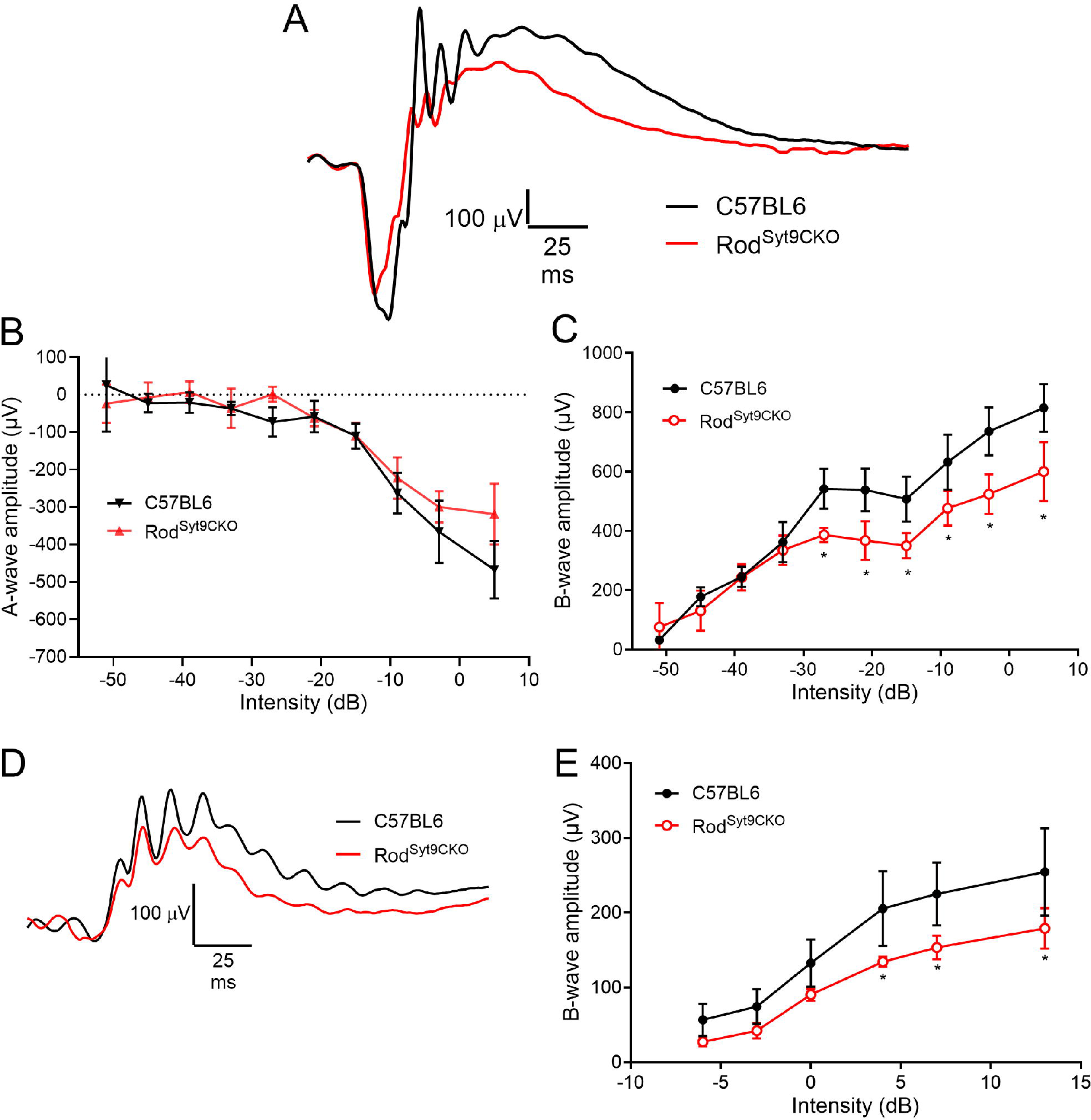
Elimination of Syt9 selectively from rods decreased scotopic and photopic ERG b-waves. (A). Representative scotopic ERG waveforms (5 dB) for control and Rod^Syt9CKO^ (n=4) mice are shown in black and red traces, respectively. (B). Scotopic a-wave amplitude as a function of flash intensity. (C). Plot of scotopic b-wave amplitude as a function of intensity. (D). Example photopic waveforms evoked from a high intensity flash (13 dB). (E). Plot of photopic b-wave amplitude as a function of flash intensity. *, P<0.01. Error bars show + S.D.

### ERG effects of selectively eliminating Syt9 from rods

Fig. 4A shows representative scotopic ERGs evoked by a high intensity 20 ms flash applied to dark-adapted control mice and Rod^Syt9CKO^ mice that lack Syt9 in rods. Scotopic a- and b-wave amplitudes are plotted in Fig. 4B and C, respectively. Selective elimination of Syt9 from rods in Rod^Syt9CKO^ mice reduced scotopic b-waves by up to 26% without significant alterations in the a-wave. Photopic ERG example waveforms are shown in Fig. 4D. Surprisingly, although photopic b-waves were not affected by loss of Syt9 from cones, they were diminished by up to 30% following selective elimination of Syt9 from rods in Rod^Syt9CKO^ mice (Fig. 4E).

### Oscillatory potentials

We next looked at effects of Syt9 deletion on ERG oscillatory potentials (OPs) that arise from interactions with amacrine cells (Liao et al., 2023; Wachtmeister, 1998). Fig. 5 displays measurements of OPs from Syt9 mutant mice under scotopic and photopic conditions. A, A’, and A’’ show representative OPs from CMV^Syt9^, Cone^Syt9CKO^, and Rod^Syt9CKO^ mice, respectively. Fig. 5B, B’, and B’’ plot scotopic OPs for the five highest intensity flashes of the scotopic ERG protocol. Photopic OPs are plotted in Fig. 5C, C’, and C’’. The only significant change in OP amplitudes in CMV^Syt9^ mice was an increase in scotopic OPs at the brightest flash intensity (Fig. 5B; t-test with Holm-Sidak correction for multiple comparisons, P = 0.00142). Eliminating Syt9 from cones had no effect on OPs under scotopic or photopic conditions (Fig. 5B’, 5C’). However, eliminating Syt9 from rods reduced OP amplitudes under scotopic conditions, with a 50% reduction at the highest flash intensity (Fig. 5B’’; P < 0.00005), but only small changes to OPs under photopic conditions (Fig. 5C’’).

**Fig. 5.**
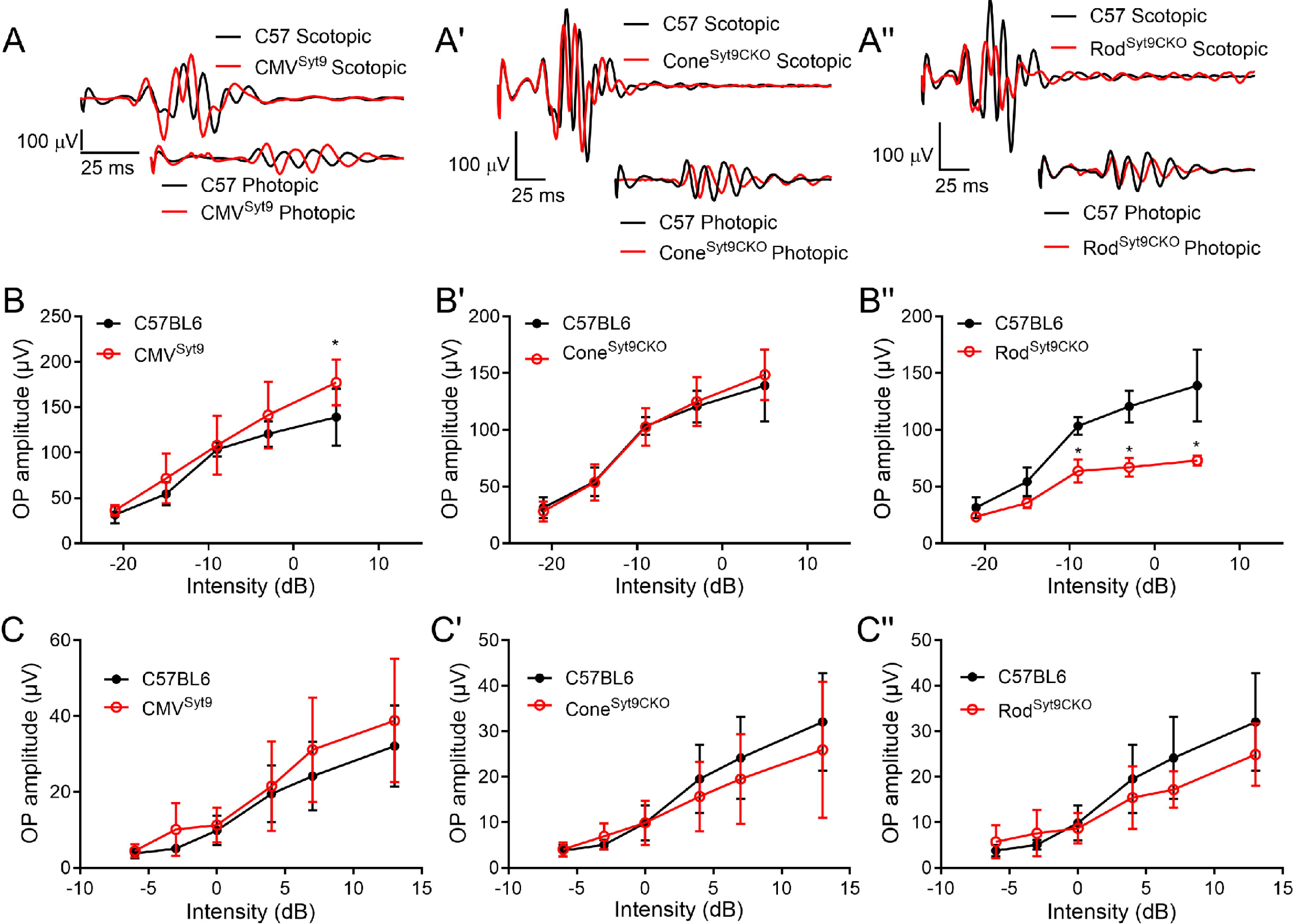
Effects of Syt9 elimination from the whole retina on ERG oscillatory potentials. (A, A’, A’’). Scotopic and photopic OPs extracted from representative ERG waveforms (5 dB and 13 dB, respectively) by bandpass filtering to remove frequencies below 70 Hz and above 280 Hz in CMV^Syt9^, Cone^Syt9CKO^, and Rod^Syt9CKO^ mice, respectively (B, B’, B’’). Scotopic OPs in CMV^Syt9^, Cone^Syt9CKO^, and Rod^Syt9CKO^ mice, respectively (C, C’, C’’). *, P<0.01. Error bars show + S.D.

### Whole cell recordings from rods lacking Syt9

Reductions in b-wave amplitude without accompanying changes in a-waves are generally evidence of impaired synaptic release from photoreceptors. Given that selective deletion of Syt9 from rods reduced scotopic b-waves at high intensities (Fig. 4), we recorded directly from rods lacking Syt9 (using CMV^Syt9^ mice). To assess release, we measured anion currents activated in rods by binding of glutamate to presynaptic EAAT5 glutamate transporters (Grassmeyer et al., 2019). The whole cell pipette solution contained SCN^-^ as the principal anion to enhance glutamate transporter anion currents (I_A(glu)_). We measured inward currents immediately after termination of depolarizing steps from -70 to -10 for 5, 25, and 500 ms. Fig. 6A shows example waveforms of I_A(glu)_ evoked by 25 ms steps in control (top black trace) and Syt9 KO rods (bottom red trace). Passive capacitative and resistive currents were subtracted using P/6 protocols. We measured both amplitude and charge transfer of I_A(glu)._ As illustrated in Figs. 6B and C, we saw no differences in evoked glutamate release between rods lacking Syt9 and a sample of control rods from C57Bl6J mice with either measurement (Fig. 6). We also analyzed spontaneous release in rods with and without Syt9. The downward transients in the example I_A(glu)_ waveforms in Fig. 6D each represent release of a single synaptic vesicle. Spontaneous release was assessed using 30 s trials with rods held at -70 mV. Spontaneous release of glutamate-filled vesicles persisted in the absence of Syt9 with no change in event frequency (Syt9KO: 0.637 + 0.486 v/s, n=10 rods; Control: 0.641 + 0.644 v/s, n=13 rods; nested t-test, P=0.85).

**Fig. 6.**
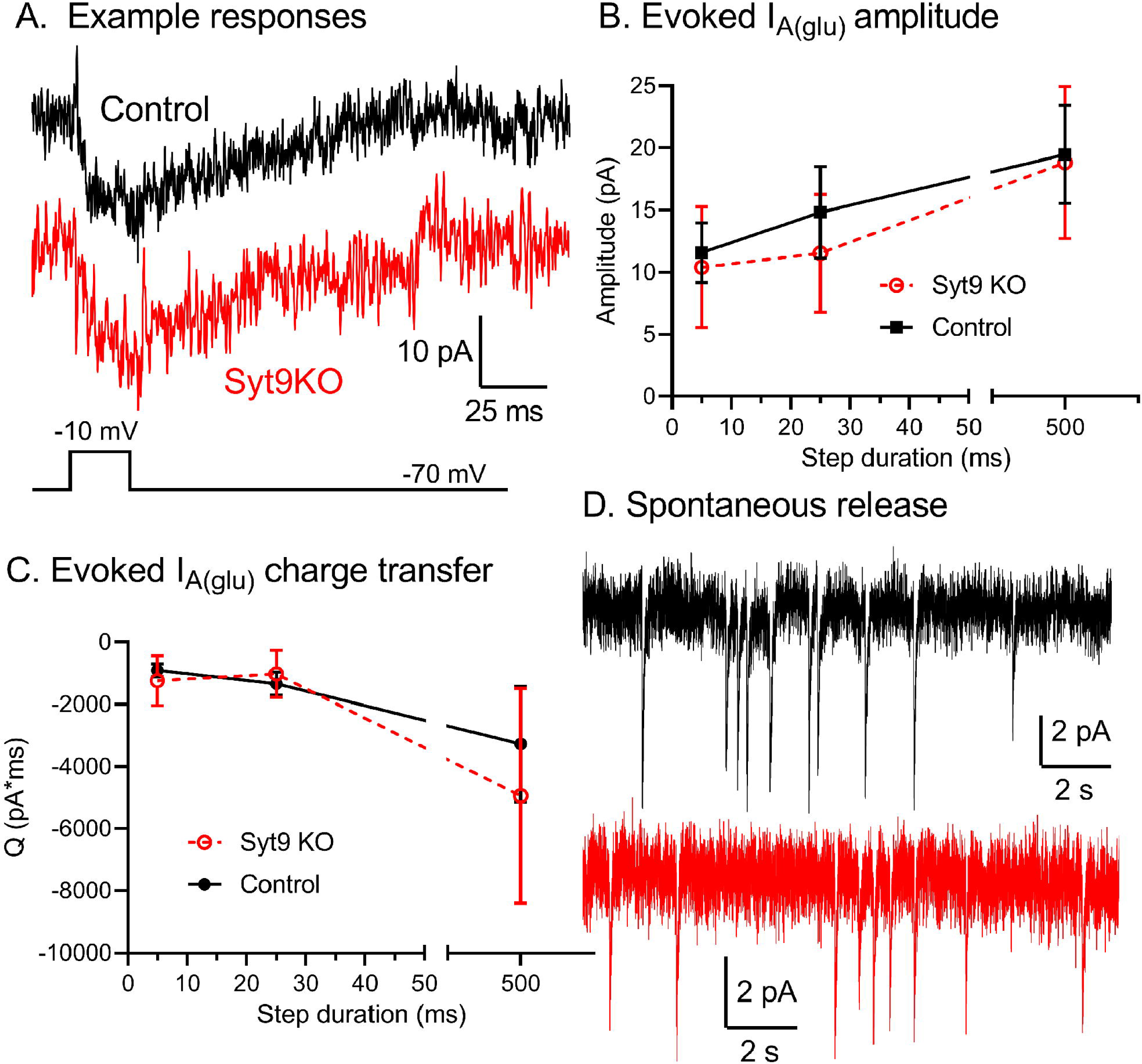
Whole cell recordings of glutamate transporter anion currents (I_A(glu)_) revealed no significant differences between control and Syt9 KO rods. A. Example I_A(glu)_ recordings from control and Syt9 KO rods evoked by 25 ms steps from -70 to -10 mV. Passive capacitative and resistive currents were subtracted using P/6 protocols. B. I_A(glu)_ amplitude from control (5 ms, n=34; 25 ms, n=27; 500 ms, n=24) and Syt9 KO rods (5 ms, n=15; 25 ms, n=16; 500 ms, n=11). Control data from (Mesnard et al., 2022). Response amplitude was measured 2 ms after termination of the test step. C. I_A(glu)_ charge transfer from control (5 ms, n=37 rods; 25 ms, n=26; 500 ms, n=20; n=15 mice) vs. Syt9 KO rods (5 ms, n=15; 25 ms, n=16; 500 ms, n=11; 4 mice). Charge transfer was measured beginning 2 ms after termination of the test step until current recovered to baseline. Baseline was defined as the current level at the end of the 2 s trial. D. Example recordings of spontaneous I_A(glu)_ release events showing release of individual glutamate-filled synaptic vesicles. Rods were held at -70 mV when measuring spontaneous release. Error bars show 95% confidence intervals.

## Discussion

Global Syt9 deletion had different effects on the ERG than selective deletion from rods or cones in mouse retina suggesting involvement of Syt9 at multiple sites in the retina. Global elimination of Syt9 from retina *increased* the amplitude of scotopic ERG b-wave responses, while deletion from rods alone *decreased* scotopic ERG b-waves and OPs. Selective deletion of Syt9 from cones had no observable effect.

Consistent with our RNAscope data showing stronger expression of Syt9 mRNA in the inner nuclear layer than in photoreceptors, RNAseq data also shows higher Syt9 transcript levels in bipolar cells compared to photoreceptors (Hartl et al., 2017; Sarin et al., 2018).

Selective deletion of Syt9 from rods showed effects only at higher flash intensities where cones contribute more significantly. However, deleting Syt9 from cones had no effect. Furthermore, deleting Syt9 from rods reduced photopic b-waves even though photopic responses are driven entirely by cones (Ronning et al., 2018) and eliminating Syt1 from cones is sufficient to abolish photopic ERG responses (Grassmeyer et al., 2019). Rods and cones are coupled to one another via Cx36 gap junctions allowing rod signals to enter cone terminals where they can be transmitted to post-synaptic neurons (Fain and Sampath, 2018). Passage of signals the other way has not been carefully investigated but one possible explanation for these effects is that loss of Syt9 in rods somehow restricts the ability of rods to convey cone signals to downstream neurons. Since release from rods themselves is unimpaired, loss of Syt9 may somehow interfere with gap junction function, perhaps by altering protein trafficking as seen with other Syt isoforms (Wolfes and Dean, 2020). Although we saw no gross anatomical changes, another possible explanation is that the loss of Syt9 from rods or other neurons may alter synaptic connectivity in subtle ways.

GABA and glycine are the major inhibitory neurotransmitter in the vertebrate retina (Eggers and Lukasiewicz, 2011) and Syt9 has been shown to modulate GABA release in cultured striatal neurons (Xu et al., 2007). These results led us to consider the possibility that loss of Syt9 might also interfere with GABA and/or glycine release from retinal neurons. Like effects of global deletion of Syt9, blocking GABA_A_ receptors with bicuculline typically enhances ERG b-waves (Arnarsson and Eysteinsson, 1997; Gottlob et al., 1988; Hirasawa et al., 2021; Vitanova et al., 2001). By contrast, loss of GABA_C_ receptors and application of GABA_C_ antagonists have been found to reduce ERG b-waves (Kapousta-Bruneau, 2000; Moller and Eysteinsson, 2003; Wang et al., 2015). This suggests that if Syt9 is modulating GABA release, it is likely to be acting predominantly at synapses possessing GABA_A_ receptors. Blocking glycine receptors with strychnine can also increase ERG b-waves (Arnarsson and Eysteinsson, 1997; Popova, 2000), although this has not been seen in every preparation (Hirasawa et al., 2021).

The OPs that ride on top of the ERG b-wave appear to arise from reciprocal synapses between rod bipolar cells and AII and A17 amacrine cells in the inner retina (Liao et al., 2023; Wachtmeister, 1998). Loss of Syt9 from rods led to a reduction in scotopic OPs that paralleled reductions in scotopic b-waves. This suggests that diminished rod input accompanying loss of Syt9 led in turn to diminished amacrine cell activity. Global deletion of Syt9 had smaller effects on OPs, with a small increase in scotopic OPs at the highest intensities. The reduction in OPs produced by eliminating Syt9 from rods might be countered by increased amacrine cell activity mediated by actions of Syt9 in other cell types. Consistent with a contribution to shaping OPs from diminished GABA release in Syt9 global knockout mice, reducing GABAergic inhibition in GABA_C_ receptor knockouts can also enhance OPs (McCall et al., 2002). Further studies on the sites and roles of Syt9 in the retina are needed to distinguish these various possibilities.

